# Statistical significance in DTI group analyses: How the choice of the estimator can inflate effect sizes

**DOI:** 10.1101/755140

**Authors:** Szabolcs David, Hamed Y. Mesri, Max A. Viergever, Alexander Leemans

## Abstract

Diffusion magnetic resonance imaging (dMRI) is one of the most prevalent methods to investigate the micro- and macrostructure of the human brain *in vivo*. Prior to any group analysis, dMRI data are generally processed to alleviate adverse effects of known artefacts such as signal drift, data noise and outliers, subject motion, and geometric distortions. These dMRI data processing steps are often combined in automated pipelines, such as the one of the Human Connectome Project (HCP). While improving the performance of processing tools has clearly shown its benefits at each individual step along the pipeline, it remains unclear whether – and to what degree – choices for specific user-defined parameter settings can affect the final outcome of group analyses. In this work, we demonstrate how making such a choice for a particular processing step of the pipeline drives the final outcome of a group study. More specifically, we performed a dMRI group analysis on gender using HCP data sets and compared the results obtained with two diffusion tensor imaging estimation methods: the widely used ordinary linear least squares (OLLS) and the more reliable iterative weighted linear least squares (IWLLS). Our results show that the effect sizes for group analyses are significantly smaller with IWLLS than with OLLS. While previous literature has demonstrated higher estimation reliability with IWLLS than with OLLS using simulations, this work now also shows how OLLS can produce a larger number of false positives than IWLLS in a typical group study. We therefore highly recommend using the IWLLS method. By raising awareness of how the choice of estimator can artificially inflate effect size and thus alter the final outcome, this work may contribute to improvement of the reliability and validity of dMRI group studies.

## 1 Introduction

Diffusion magnetic resonance imaging (dMRI) has been used extensively to study fundamental biological concepts (Assaf et al., 2019; Novikov et al., 2019), pathologies of the brain (Cercignani and Gandini Wheeler-Kingshott, 2019; Lunven et al., 2015; Phillips et al., 2016; Sabia et al., 2017), and the architectural configuration of white matter (WM) tracts (Catani et al., 2013; David et al., 2019; Thiebaut de Schotten et al., 2012). As dMRI became more commonly used, there was a need to improve its reliability for clinical applications (Eierud et al., 2014; Nir et al., 2013; Owen et al., 2013; Rudie et al., 2013; Schwarz et al., 2013). Methodological developments that contributed to this improvement are related to cardiac gating (Chang et al., 2005; Kozák et al., 2013), high-field MRI scanners (Moser et al., 2017), stronger and faster switching MR gradients (McNab et al., 2013; Setsompop et al., 2013), image reconstruction techniques (Lustig et al., 2007), diffusion model estimation approaches (Collier et al., 2018; Pannek et al., 2012; Tax et al., 2015; Veraart et al., 2013b), correction strategies for Gibbs-ringing (Kellner et al., 2016; Perrone et al., 2015; Veraart et al., 2016a), signal drift (Vos et al., 2017), thermal noise (St-Jean et al., 2016; Veraart et al., 2016b) eddy current distortions (Andersson et al., 2016; Andersson and Sotiropoulos, 2016, 2015), and susceptibility induced deformations (Andersson et al., 2018, 2003; Graham et al., 2017), among others.

Processing tools are the key contributors in minimizing adverse effects of confounding factors on the final results. Despite the theoretical benefits of integrating novel methodological developments in the dMRI processing pipeline, there is no consensus on which settings or algorithms should be preferred for, for instance, a typical diffusion tensor imaging (DTI) study in which two groups of subjects (e.g., healthy controls vs. patients) are compared. This lack of agreement is reinforced by our limited understanding of whether a specific processing method has a significant contribution to the reliability of the subsequent group analysis in terms of outcome. In this context, one could state that, in practice, the added benefit of a particular data correction procedure is nullified if there are other data aspects with a much higher variability. As an example, the decrease in diffusion parameter estimation bias due to Gibbs ringing correction may be completely swamped by the high noise levels in low-SNR dMRI data, obviating the relevance of performing this processing step.

In general, the relative improvement of one processing step not only depends on the intrinsic quality of the data, but also on the performance of the other processing steps used in the dMRI pipeline. Correcting spatial misalignment across multiple diffusion-weighted images (DWIs) due to subject motion, for instance, may benefit from preceding denoising of these images. In addition, after the data has been corrected for artifacts, strategies to further analyze the data (e.g., using fiber tractography, histograms, ROIs, voxel-based approaches, or network graphs) may have a difference in sensitivity to the benefit of some of the individual processing steps and potentially generate differences in the final outcome of a group study.

While many steps in a dMRI processing pipeline can be considered as optional, for several diffusion approaches such as DTI or diffusion kurtosis imaging (DKI), there is the mandatory step of choosing the diffusion estimation method to obtain model parameters. Over the last decade, a plethora of such estimators have been used, including ordinary linear least squares (OLLS), non-linear least squares (NLLS), weighted linear least squares (WLLS), and their constrained, robust and conditional extensions, among others (Andersson, 2008; Chang et al., 2012, 2005; Collier et al., 2015; Jones and Basser, 2004; Koay et al., 2009; Kristoffersen, 2012, 2007; Salvador et al., 2005; Tax et al., 2015; Veraart et al., 2013b, 2011). Assuming that data outliers have been identified and removed, a specific version of the WLLS, iterative WLLS (IWLLS), shows high performance characteristics in terms of accuracy and precision and may even be preferred over advanced NLLS estimation methods (Veraart et al., 2013b). Yet, OLLS is still the most widely used estimation method and often defined as the default in common software tools (e.g., FSL – (Jenkinson et al., 2012)).

Similar to the other dMRI processing steps, one can also question the relevance of choosing a particular diffusion estimation approach. Does it really matter which estimator is used for the final outcome of a group study? In this work, we address this concern. More specifically, we performed a dMRI group analysis using Human Connectome Project (HCP) data sets and compared the results obtained with OLLS and IWLLS. To this end, and without loss of generality, we investigated gender related differences (Caeyenberghs and Leemans, 2014; Herting et al., 2012; Hsu et al., 2008; Ingalhalikar et al., 2014; Kanaan et al., 2012; Menzler et al., 2011; Núñez et al., 2017; Tyan et al., 2017; Westerhausen et al., 2003; Wierenga et al., 2017) to evaluate the potential differences in final outcomes using the two estimators. Preliminary results of this work were presented at the International Society for Magnetic Resonance in Medicine (ISMRM) meeting in Toronto, Canada (David et al., 2015).

## 2 Methods

### 2.1 Subject data and processing

Minimally preprocessed DWIs were collected from the HCP S500 release (Essen et al., 2012; Glasser et al., 2013). Briefly, the data consist six separate acquisitions of 90 DWIs acquired with diffusion weightings (b-values) equal to 1000/2000/3000 s/mm^2^ and five, six or seven non-DWIs (b-value = 0 s/mm^2^). Every image was acquired with both left-to-right and right-to-left phase encoding directions; the voxel size was 1.25 mm isotropic. Susceptibility artifacts, eddy current induced distortions, and subject motion were corrected with the *FSL* tools taking into account any reorientations of the diffusion gradient orientations (Andersson et al., 2003; Jenkinson et al., 2012; Leemans and Jones, 2009; Sotiropoulos et al., 2013). All datasets were further processed with *ExploreDTI* version 4.8.6. (Leemans et al., 2009) using two different tensor estimation approaches: (a) OLLS (Basser et al., 1994) and (b) IWLLS (Veraart et al., 2013b). For this step, only the 90 DWIs with b-value of 1000 s/mm^2^ and 9 non-DWIs per participant were selected for diffusion tensor estimation. In addition, we also corrected for the gradient nonlinearities in the diffusion-weighted gradients during this estimation procedure (Bammer et al., 2003; Mesri et al., 2019; Sotiropoulos et al., 2013). Every participant for which all the 90 b = 1000 s/mm^2^ images were available, and which was not listed among the participants with known anatomical anomalies or data quality issues, was included in the analysis. The complete list of the excluded participants can be found on the appropriate HCP wiki page (HCP, 2017). The final sample size is 409 participants, consisting of 244 females and 165 males.

### 2.2 Voxel-based analysis

For each subject, fractional anisotropy (FA) maps were calculated from the fitted tensors (using OLLS and IWLLS) and transformed to the Montreal Neurological Institute (MNI) template via the native-to-MNI warp files, provided by the HCP team (Fonov et al., 2011). Voxelwise statistical comparisons of FA between the male and female groups were performed using the permutation analysis of linear models (PALM) (Holmes et al., 1996; Nichols and Holmes, 2003; Winkler et al., 2014), a Matlab based open-source toolbox, version alpha104 with 10000 permutations. For all the tests (next section), calculations are based on nonparametric permutations as this approach was proven to be more efficient in producing fewer false positives than parametric methods (Eklund et al., 2016). Significance was determined at p_corr_ < 0.05 using family-wise error rate (FWER) adjustment to correct for multiple comparisons after applying threshold-free cluster enhancement (TFCE) (Smith and Nichols, 2009). Calculation speed was accelerated using the tail approximation (Winkler et al., 2016). A Dell server with 72 Intel Xeon E7-8870 v3 @ 2.10 GHz dual cores with 1 TB RAM was used for calculations.

### 2.3 Statistical tests

#### 2.3.1 Effect of tensor estimator

For each participant, there are two FA maps: one obtained from the diffusion tensor estimated with OLLS and one with IWLLS. In order to investigate the potential differences in FA (regardless of gender) between the OLLS and IWLLS pipelines, we used a paired two-sample t-test. This procedure tests whether there is a significant effect of using a different tensor estimation method on FA, without considering if the participant is female or male.

#### 2.3.2 Effect of Gender

Differences in FA values between males and females (denoted as FA_m_ and FA_f_) were investigated using an unpaired two-sample t-test for the OLLS and IWLLS pipelines separately. A further correction was applied via the *“-corrcon*” option in PALM, which accounts for the multiple contrasts during the FWER correction.

#### 2.3.3 Pipeline dependent gender differences

To test whether gender differences depend on the tensor estimation method, we performed a two-sample t-test on the gender, where the tested variable is the difference in FA, denoted as Δ*FA*, between the IWLLS and OLLS pipelines:

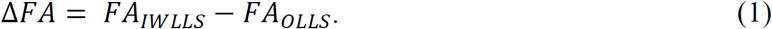

More specifically, we evaluated with this test whether the Δ*FA* values for males, denoted as Δ*FA*_m_, differ significantly from the Δ*FA* vales for females, denoted as Δ*FA*_f_. Statistically, this procedure is the same as the interaction part of a two-group analysis of variance (ANOVA) test with two levels per participant. A significant effect means that the gender differences are solely driven by the choice of estimation method. Independent and symmetric errors were assumed to boost the statistical power of the test, by using the command “*-ise*” in PALM. Effect sizes and their distributions were analyzed in detail within the regions of significance.

#### 2.3.4 Effect size

The practical significance of the findings was further evaluated by reporting effect sizes, as suggested by the American Statistical Association’s (ASA) recent statement on p-values: “*A p-value, or statistical significance, does not measure the size of an effect or the importance of a result*.” (Wasserstein and Lazar, 2016). Accordingly, we used Cohen’s *d*, a frequently applied effect size estimator. Furthermore, because Cohen’s *d* is not a robust effect size measure to outliers, skewness, heavy-tails and the combinations of these factors, the shape differences between the voxelwise distributions of FA values were studied via the shift function (Rousselet et al., 2017). The 95% percentile confidence intervals for the decile differences were estimated with a bootstrap estimation (1000 samples), using the Harrell–Davis estimator (Wilcox, 2012), as implemented in the *Matlab Robust Graphical Methods For Group Comparisons* (matrogme) toolbox, version 0.0.9000 (Rousselet et al., 2017).

## 3 Results

### 3.1 Effect of tensor estimator

Fig. 1 shows the result for the paired t-test that investigates the difference in FA between the OLLS and IWLLS estimation methods. To further emphasize the differences, we show the effect size (with Cohen’s *d*) only for the voxels that were statistically significant after applying the multiple comparison correction procedure. The map shows that these differences are significant in the whole brain and are tissue-dependent. Larger effect sizes were revealed in the core WM, such as in the corpus callosum (CC), the corticospinal tract (CST), and the optic radiation (OR), where FA values are relatively high. Areas with lower FA values near the cortical and deep GM regions (thalamus, hippocampus, putamen, etc.) resulted in no or negligible differences, as expressed by the white areas in the image that indicate a near zero effect size. Overall, the IWLLS estimator results in significantly higher FA values in the vast majority of the WM compared to using OLLS.

**Fig. 1.**
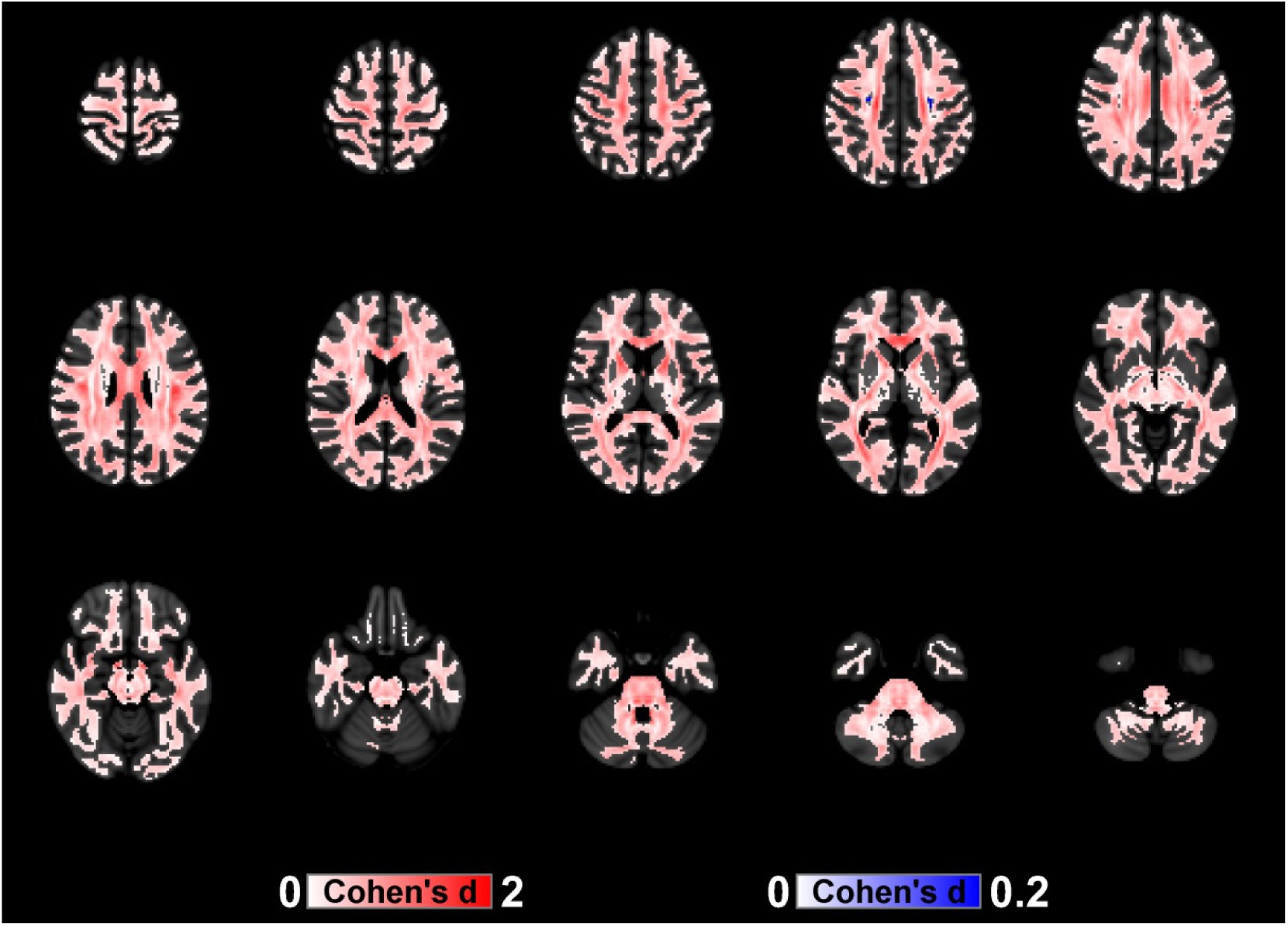
Effect sizes (defined as Cohen’s *d*) are shown as color maps overlaid on regions with statistically significant differences in FA between using the IWLLS and OLLS estimators, presented in MNI space. Notice the different color scale magnitudes for the effect sizes. The reddish and blueish color bars reflect regions where Δ*FA* > 0 and Δ*FA* < 0, respectively (see Eq. 1). (Radiological view: left on the image is right in the brain and vice versa).

The systematic deviation in FA between OLLS and IWLLS is further highlighted in Fig. 2, where the FA values are averaged across all 409 subjects. It is clear that for most of the WM voxels (∼FA>0.2) the mean FA values are higher for the IWLLS estimator than for the OLLS estimator.

**Fig. 2.**
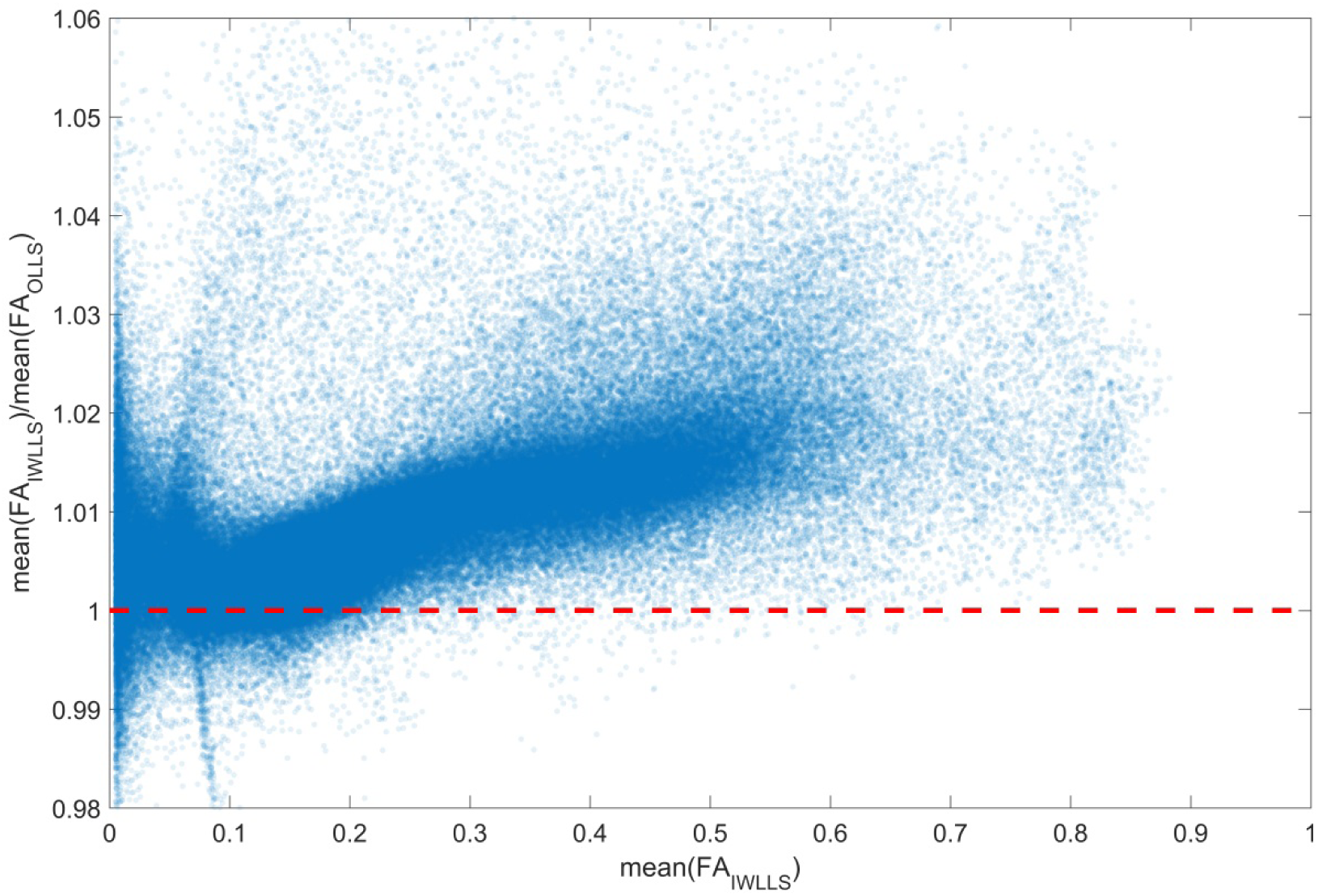
Scatterplot of the ratios of the FA values from the IWLLS and OLLS estimators as a function of FA from the IWLLS estimator. Each point in the scatterplot represents the average FA value across all 409 subjects for each brain voxel in MNI space. If there was no systematic deviation between the OLLS and IWLLS estimators, the points should be located around the unity value, indicated by the red dashed line.

### 3.2 Effect of gender

Fig. 3 shows the result of the voxelwise two-sample t-tests for both the OLLS and the IWLLS estimator, indicating the regions where FA_f_ > FA_m_ with p_corr_ < 0.05. The results of the opposite tests, that is, the regions where FA_m_ > FA_f_ with p_corr_ < 0.05, are shown for both OLLS and IWLLS in Suppl. Fig. 1. Note that the overlap itself of the two tests does not necessarily indicate identical results. In addition, the lack of overlap is not indicative of a difference in outcome between the OLLS and IWLLS results. At this stage, the results merely illustrate that there is general agreement in spatial overlap of the regions that were deemed significant in terms of FA based gender differences.

**Fig. 3.**
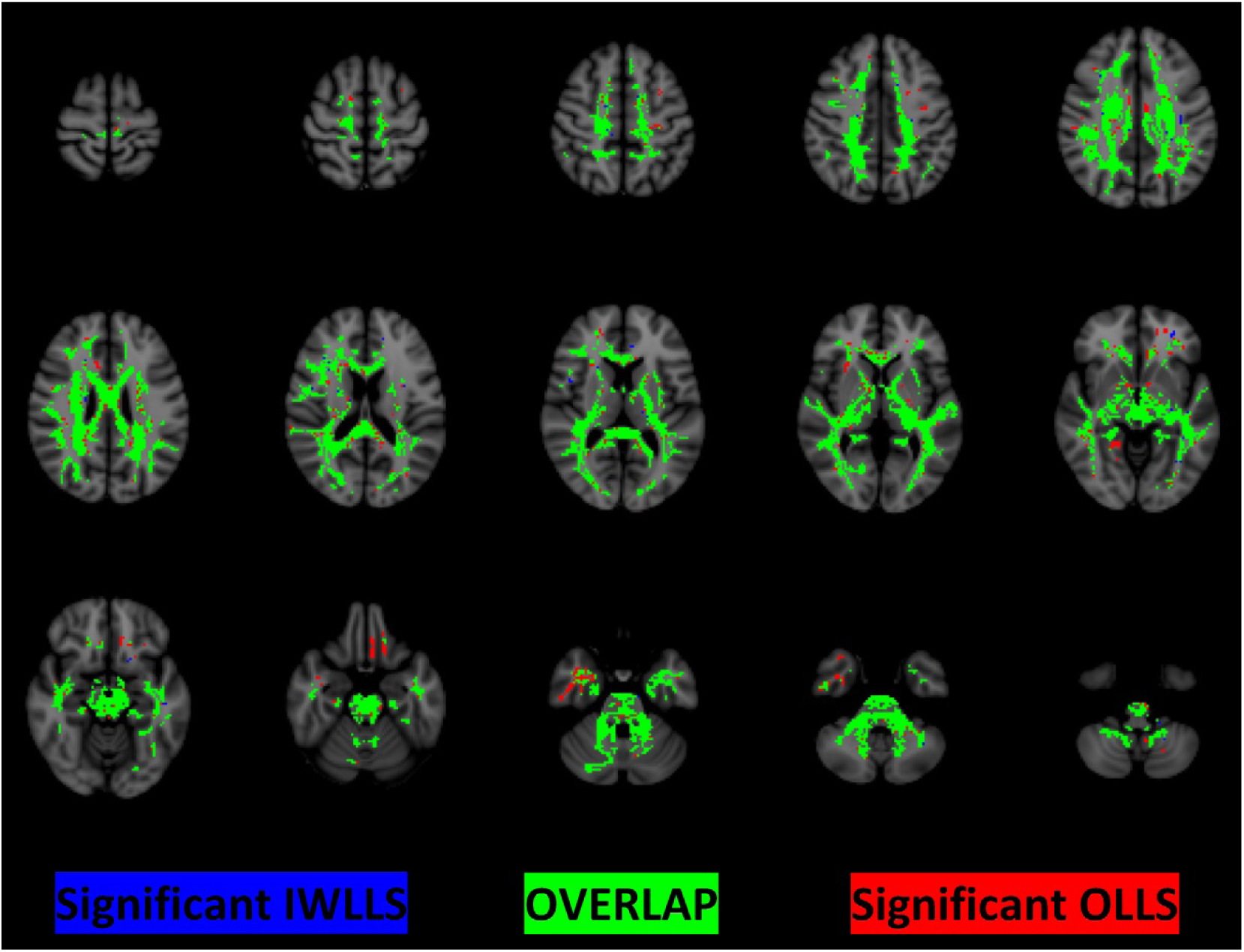
Results of the voxelwise analysis, indicating the regions where FA is significantly higher for females than males. Voxels colored in red and blue represent the regions where FA estimates were obtained with OLLS and IWLLS, respectively. The green voxels show their overlap, i.e., the regions where both OLLS and IWLLS reflect significantly higher FA values for females than for males. (Radiological view: left on the image is right in the brain and vice versa).

### 3.3 Pipeline dependent gender differences

Fig. 4 shows to which extent gender-based FA differences are driven by the choice of estimator (i.e., using OLLS or IWLLS). Overall, gender differences depend on the choice of estimator mainly in the following areas with p_corr_ < 0.05: parts of the CC and brainstem for ΔFA_m_ > ΔFA_f_ and parts of the CST for ΔFA_f_ > ΔFA_m_. To get a more detailed insight into the effect of estimation choice on the observed gender-based FA differences, we investigate the four possible scenarios (FA_f_ > FA_m_ or FA_m_ > FA_f_ in regions where ΔFA_m_ > ΔFA_f_ or ΔFA_f_ > ΔFA_m_) in the following subsections.

**Fig. 4.**
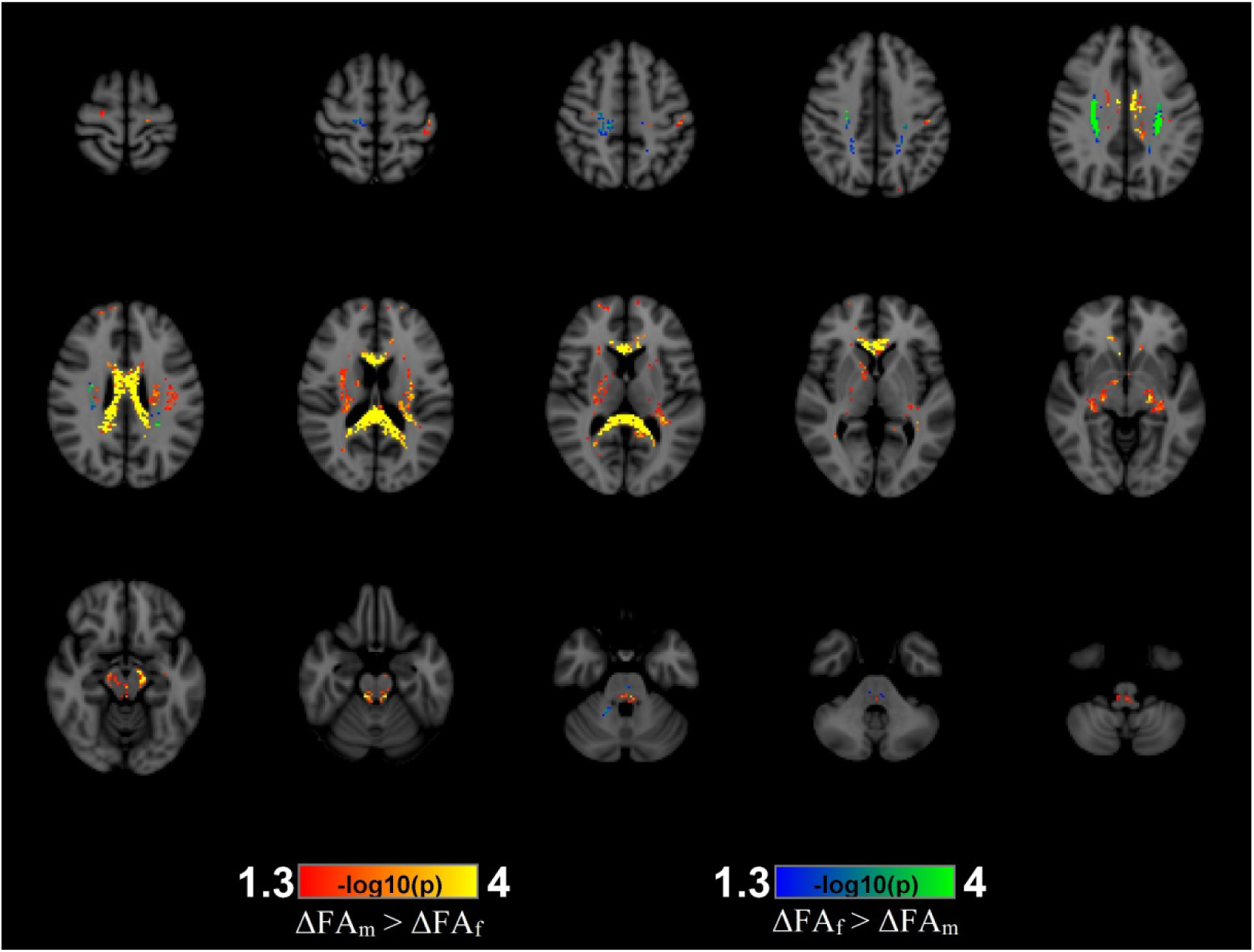
Significance maps are shown for the interaction of estimator choice with gender-based FA differences. To enhance the contrast for significance, color-encoding is according to -log_10_(p-value) with minimum and maximum values of -log_10_(0.05) ≈ 1.3 and -log_10_(1/10000) = 4 (1/10000 is the smallest achievable p-value with 10000 permutations), respectively. The difference in color encoding reflects how the choice of estimator can drive the gender-based FA difference in opposite directions, i.e., ΔFA_m_ > ΔFA_f_ (red-to-yellow coloring) and ΔFA_f_ > ΔFA_m_ (blue-to-green coloring). (Radiological view: left on the image is right in the brain and vice versa).

#### 3.3.1 Scenario 1: FA_f_ > FA_m_ in regions of ΔFA_m_ > ΔFA_f_

Fig. 5 a) shows the area of investigation. The generality of the estimator-induced bias can be seen on Fig. 5 b), which shows the differences of the effect sizes as a function of OLLS-based effect sizes.

**Fig. 5.**
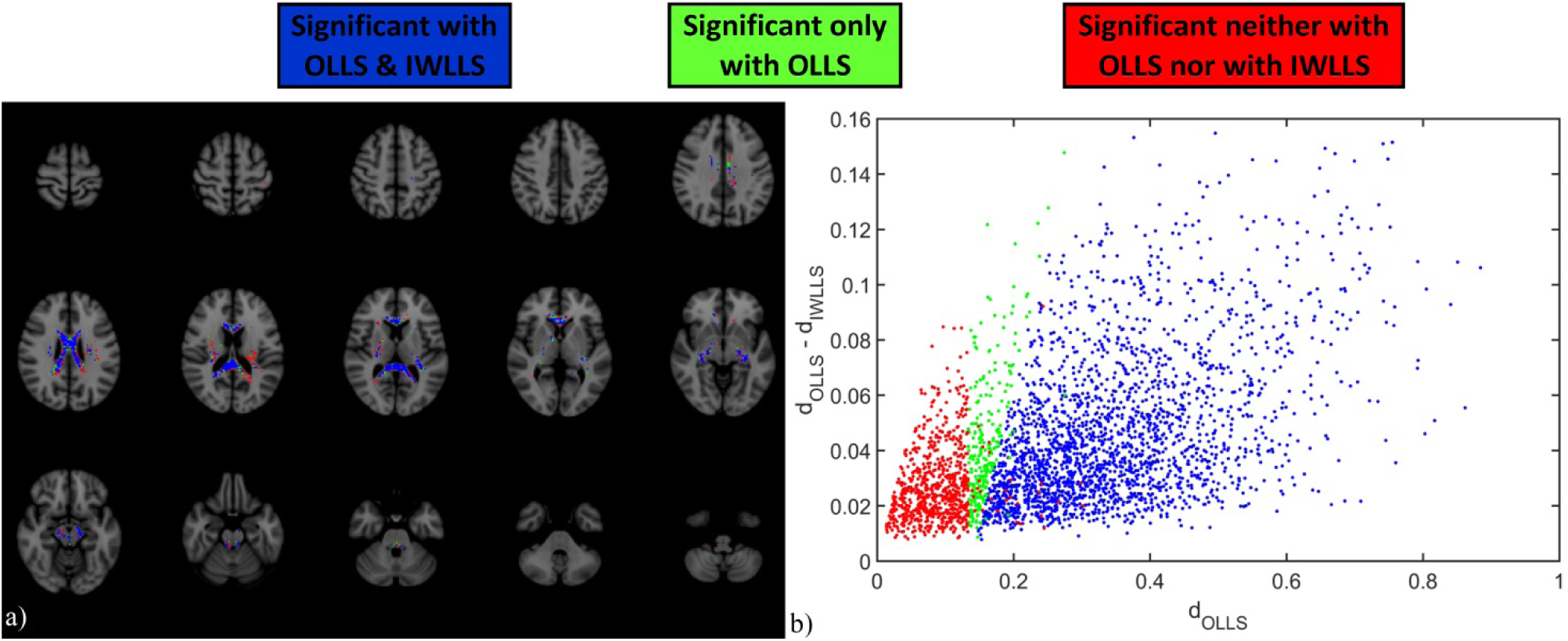
a) The spatial distribution of the voxels in MNI space, where males have a significantly larger ΔFA than females and where FA_f_ > FA_m_, regardless of whether the test was significant or not with any of the estimators. There were no voxels where the IWLLS-based FA_f_ > FA_m_ test was significant, while the OLLS-based was not. b) Scatterplot of the difference in effect sizes between OLLS (*d*_OLLS_) and IWLLS (*d*_IWLLS_) based effect sizes as a function of *d*_OLLS_. (Radiological view: left on the image is right in the brain and vice versa).

To get a better insight into the underlying effect of how estimator choice can drive gender-based FA differences, we explicitly show the data points of all participants for a single voxel. To showcase this effect, we performed a detailed analysis for the voxel in which the effect size of the FA_f_ > FA_m_ test decreased the most, when the estimation was changed from OLLS to IWLLS (Fig. 6). MNI coordinates of this voxel, located in the midsagittal plane of the splenium, are: x = 0; y = -38; z = 16. Figs. 6 a) and b) show the distribution of FA values from all subjects in the given voxel when using the OLLS (FA_OLLS_) and IWLLS (FA_IWLLS_) estimators, respectively. The effect size is lower for IWLLS than for OLLS: Cohen’s *d* decreased from 0.49 to 0.34. By investigating the FA_IWLLS_ / FA_OLLS_ ratios (Fig. 6 c)), it can be readily seen that FA_m_ increased more than FA_f_ when changing the estimator from IWLLS to OLLS. The FA_m_ - FA_f_ difference is plotted for each decile with the bootstrapped confidence intervals as a function of male deciles, indicating that the increase in FA_m_ was systematically larger than the increase in FA_f_ by 0.5-2% due to this change (Fig. 6 d)). Note that if a confidence interval does not include zero, one may also conclude that said difference is significant between the changes of these ratios.

**Fig. 6.**
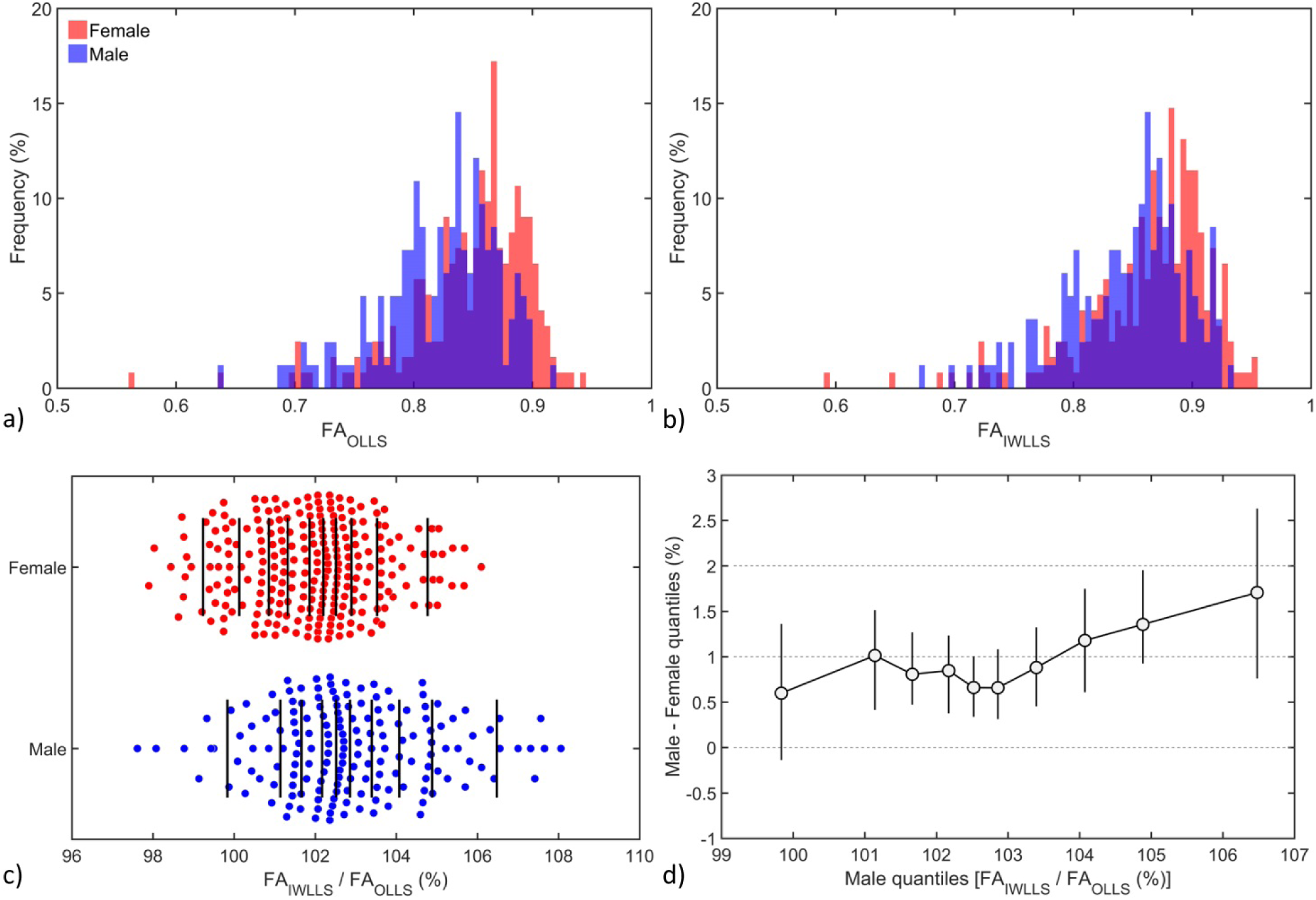
The FA distribution for males (blue) and females (red) for OLLS (a) and IWLLS (b), respectively, in a voxel located in the corpus callosum (CC), where the effect size decreased the most from *d*_OLLS_ = 0.49 to *d*_IWLLS_ = 0.34. c) The ratio of FA_IWLLS_ / FA_OLLS_ per gender, with the vertical lines indicating the deciles. d) The quantile differences between males and females for the ratios shown in panel c).

#### 3.3.2 Scenario 2: FA_f_ > FA_m_ in regions of ΔFA_f_ > ΔFA_m_

Fig. 7 a) shows the area of investigation. The generality of the estimator-induced bias can be seen on Fig. 7 b), which shows the differences of the effect sizes as a function of IWLLS-based effect sizes.

**Fig. 7.**
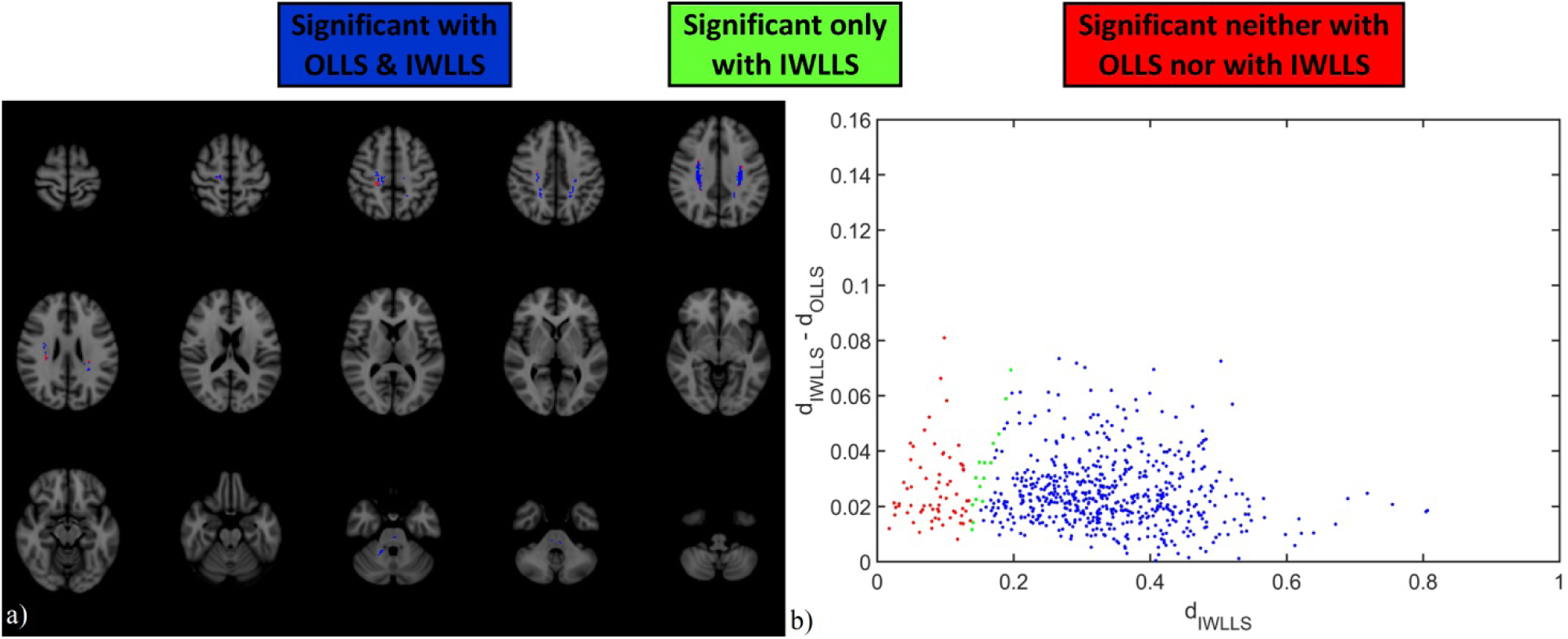
a) shows the spatial distribution of the voxels in MNI space, where females have a significantly larger ΔFA than males and where FA_f_ > FA_m_, regardless of whether the test was significant or not with any of the estimators. There were no voxels where the OLLS-based FA_f_ > FA_m_ test was significant, while the IWLLS-based was not. b) Scatterplot of the difference in effect sizes between OLLS (*d*_OLLS_) and IWLLS (*d*_IWLLS_) based effect sizes as a function of *d*_IWLLS_. (Radiological view: left on the image is right in the brain and vice versa).

Fig. 8 shows the detailed analysis for the voxel in which the effect size of the FA_f_ > FA_m_ test increased the most, when the estimation was changed from OLLS to IWLLS. MNI coordinates of the voxel, located in the superior longitudinal fasciculus (SLF), are: x = 28; y = -20; z = 36. Figs. 8 a) and b) show the distribution of FA values from all subjects in the given voxel when using the OLLS (FA_OLLS_) and IWLLS (FA_IWLLS_) estimators, respectively. The effect size is higher for IWLLS than for OLLS: Cohen’s *d* increased from 0.19 to 0.27. Fig. 8 c) shows the FA_IWLLS_ / FA_OLLS_ ratios per gender, indicating that FA_f_ increased more than FA_m_ when changing the estimator from IWLLS to OLLS. Fig. 8 d) shows the shift function. The FA_m_ - FA_f_ difference is plotted for each decile with the bootstrapped confidence intervals as a function of male deciles, indicating that the increase in FA_f_ over FA_m_ was larger with 1-2%, except in the highest decile, where FA increased nearly at the same rate. Note that if a confidence interval does not include zero, one may also conclude that said difference is significant between the changes of these ratios.

**Fig. 8.**
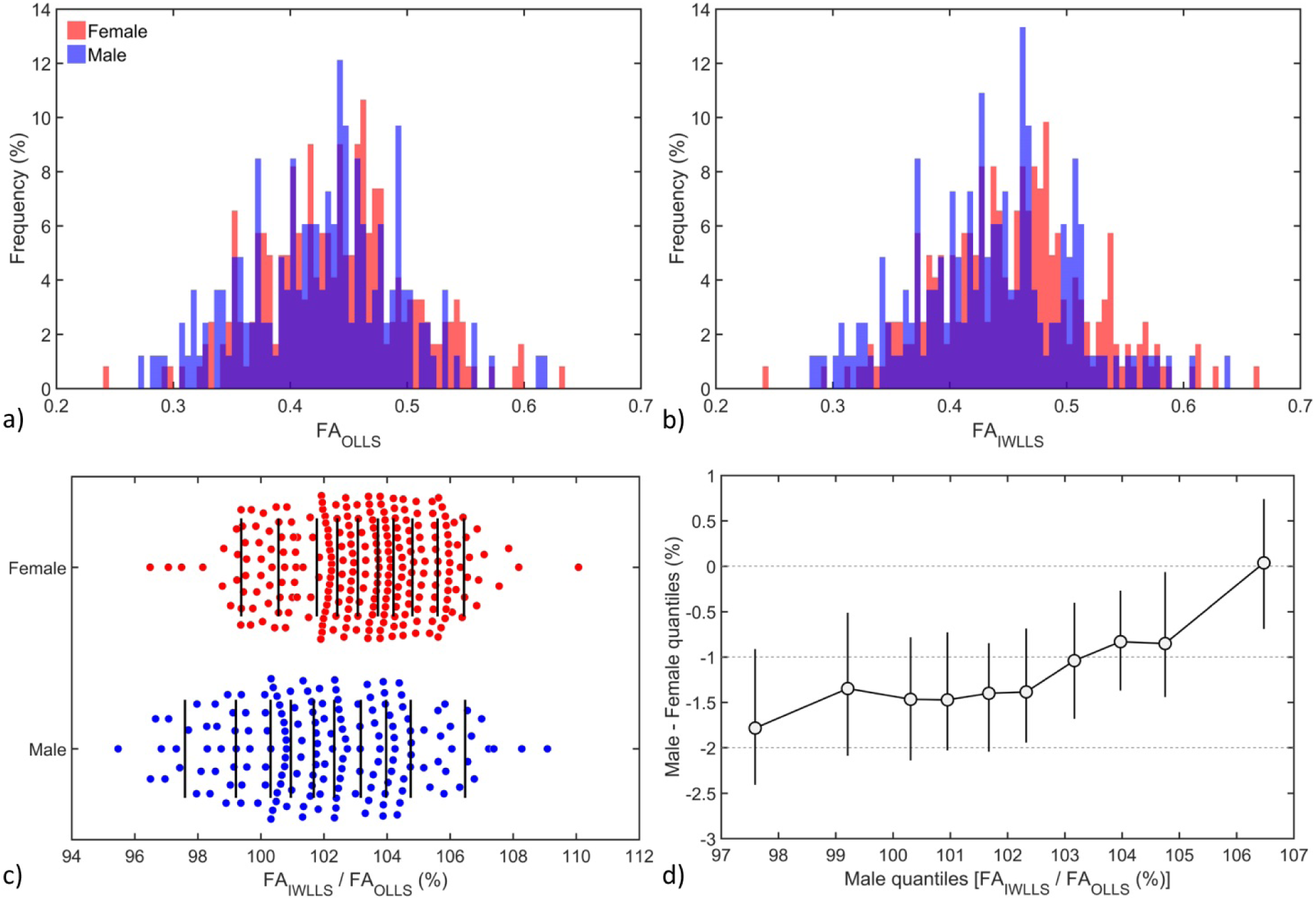
The FA distribution for males (blue) and females (red) for OLLS (a) and IWLLS (b), respectively, in a voxel located in the superior longitudinal fasciculus (SLF), where the effect size increased the most from *d*_OLLS_ = 0.19 to *d*_IWLLS_ = 0.27. c) The ratio of FA_IWLLS_ / FA_OLLS_ per gender, with the vertical lines indicating the deciles. d) The quantile differences between males and females for the ratios shown in panel c).

#### 3.3.3 Scenario 3: FA_m_ > FA_f_ in regions of ΔFA_m_ > ΔFA_f_

Males have a smaller area where FA_m_ > FA_f_, therefore the area where estimators could have any effect is also smaller compared to females. The area of investigation is located where ΔFA_m_ > ΔFA_f_ is significant, as shown in Fig. 4, but within that region it is limited to voxels where FA_m_ > FA_f_. Fig. 9 shows the differences of the effect sizes as a function of IWLLS-based effect sizes. For the sake of simplicity, the spatial distribution of the voxels in MNI space is not shown.

**Fig. 9.**
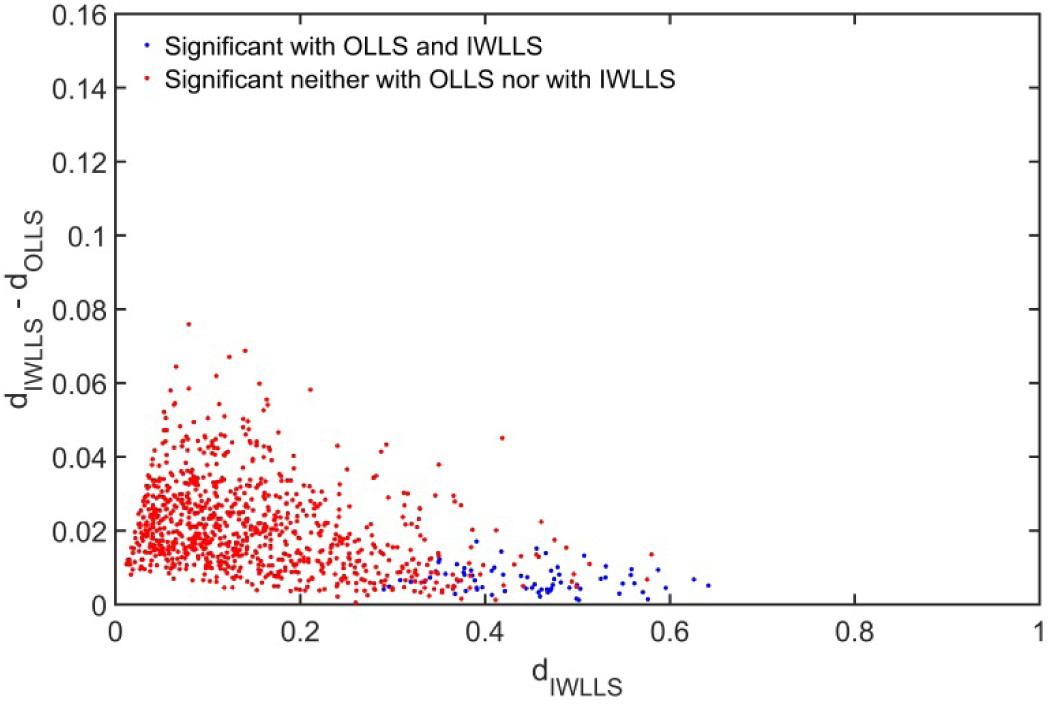
Scatterplot of the difference in effect sizes between OLLS (*d*_OLLS_) and IWLLS (*d*_IWLLS_) based effect sizes as a function of *d*_IWLLS_, where males have a significantly larger ΔFA than females and where FA_m_ > FA_f_.

#### 3.3.4 Scenario 4: FA_m_ > FA_f_ in regions of ΔFA_f_ > ΔFA_m_

The area of investigation is located where ΔFA_f_ > ΔFA_m_ is significant, as shown in Fig. 4, but within that region is limited to voxels where FA_m_ > FA_f_. Fig. 10 shows the differences of the effect sizes as a function of OLLS-based effect sizes. For the sake of simplicity, the spatial distribution of the voxels in MNI space is not shown.

**Fig. 10.**
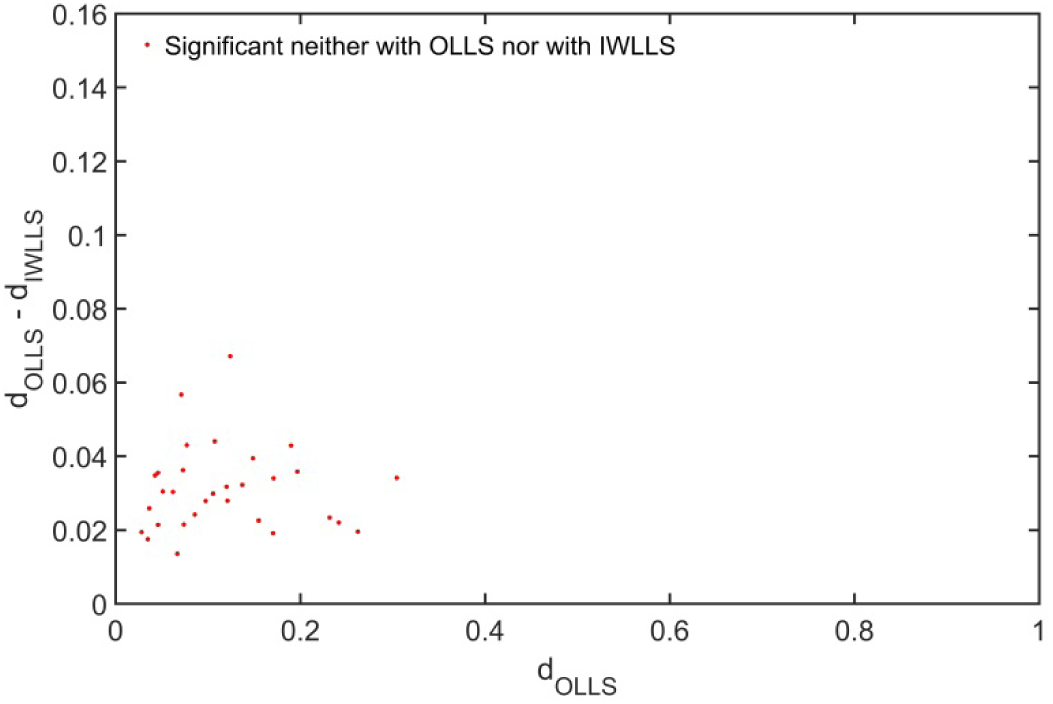
Scatterplot of the difference in effect sizes between OLLS (*d*_OLLS_) and IWLLS (*d*_IWLLS_) based effect sizes as a function of *d*_OLLS_, where females have a significantly larger ΔFA than males and where FA_m_ > FA_f_.

## 4 Discussion

In this work, we investigated how making a different choice for a specific data processing step can affect the outcome in a typical DTI group study. More specifically, we performed a voxel-based analysis, comparing FA values between males and females using HCP data, and revealed that a higher effect size was obtained with the OLLS diffusion tensor estimator than with its IWLLS counterpart. If we consider that the IWLLS estimator has a higher accuracy, we can conclude that OLLS overestimates the observed FA based gender differences. With the majority of published DTI studies having used the OLLS estimator, it is not hard to imagine that the lack of general agreement in findings for several research topics (both in neuroscience and clinical applications) could also be partly attributed to the higher number false positives introduced by the OLLS estimator as compared with the IWLLS estimator. In the following paragraphs, we will discuss how our findings relate with what is known in functional MRI (fMRI) and we will place our results in the context of other dMRI studies.

The term ‘blobology’ (Poldrack, 2012) corresponds to the colorful patches, the ‘blobs’, of fMRI brain studies, summarizing the localization of the results after processing and statistical thresholding. The phrase reflects an inherent frustration within the neuroimaging community, partly due to the lack of effect size reports. In dMRI studies, unfortunately, effect sizes are rarely reported. Researchers often spend most of their efforts on reporting statistically significant results from the data, while the extent of these effects, which is highly complementary, is hardly considered.

With large databases like ADNI (n > 2000) (Mueller et al., 2005), ENIGMA (n > 10000) (Thompson et al., 2014), HCP (n = 1200), UK BioBank (final n = 100000) (Sudlow et al., 2015), or the Whitehall study (n = 6035) (Filippini et al., 2014), the challenges are shifting toward huge sample sizes to allow the detection of small effects, which otherwise could not be identified (Smith and Nichols, 2018). But even for group studies based on these cohorts, not properly processing the data according to best practices may still result in biases that will affect the reliability of the final outcome measures. Thompson et al. (Thompson et al., 2016) reached a similar conclusion in relation to the genome-connectome association in the ENIGMA project: *“… Clearly, the ability to pursue such an approach on a large scale, within ENIGMA, depends on several factors: a working group, ENIGMA-DTI, was set up to assess its feasibility. First, unless diffusion-weighted MRI measures show greater genetic effect sizes than other traits assessed so far, there must be tens of thousands of DTI scans available from people with GWAS for such a study to be well powered …*”. According to the ENIGMA-DTI processing protocol (ENIGMA DTI protocol, 2018), the OLLS estimator is used via the *FSL* toolbox *dtifit*. In all of the aforementioned large-scale cohorts (ADNI, HCP, UK BioBank, Whitehall study), OLLS is also used which, in light of our findings, may adversely affect the reliability of the final outcome in a group study. Generally, lower-quality dMRI data in terms of effective SNR or CNR benefit more from using an estimator with better performance characteristics such as the IWLLS approach (Veraart et al., 2013a, 2013b). In this work, we used HCP data, which are among the highest quality data available in current large-scale cohorts (Bastiani et al., 2019). Given the lower number of DWIs, the lower SNR and CNR, and the higher amount of physiological artifacts in more conventional neuroimaging studies, especially in a clinical setting, one can expect even more inflated effect sizes by using the OLLS estimator than those observed in this work.

In this work, we carried out the voxelwise analysis with the Statistical Parametric Mapping (SPM) toolbox (Penny et al., 2007), rather than with another common approach, i.e. tract-based spatial statistics (TBSS) (Smith et al., 2006). While our results in this manuscript would be conceptually the same when using TBSS, confounds may arise from the skeletonization step, which may be different between the OLLS and the IWLLS. Differences in their local FA maxima could then affect statistical analysis and may further complicate interpretation of the outcome (Bach et al., 2014). Assuming the same skeleton could be provided for both datasets, e.g., via the overlap and the fusion of the skeletons, there is no reason to consider that the results presented in this work would be significantly different.

Researchers often justify the choices made for specific processing steps in their data processing pipeline by referring to previously peer-reviewed studies, which used the same settings or algorithms, despite the availability of more reliable alternatives. In addition, as OLLS generates an artificially higher effect size than IWLLS, it stimulates the positive bias in publications (Rothstein et al., 2006) and contributes to “the natural selection of bad science” (Smaldino and McElreath, 2016). To some extent, following the implementation of “registered reports” may mitigate this concern as the processing pipeline can be reviewed and scrutinized before starting the actual analysis (Nosek and Lakens, 2014).

In a recent review paper by Poldrack et al. (Poldrack et al., 2017) the lack of common consensus in processing and analysis was showcased for fMRI. With common fMRI software packages, it was shown that the number of possible analysis workflows can be as much as 69,120. For DTI, it is not hard to achieve the same order of magnitude for this number of workflows given the vast amount of options and parameter settings one can think of. In this work, we specifically investigated the effect of choosing between the OLLS and the IWLLS estimator on the outcome of the analysis, as using a diffusion tensor estimator is mandatory. Other processing steps, such as denoising and correcting for artifacts are not per se necessary (although highly recommended, of course) to continue with performing an actual group study. In this context, there may be several aspects of a typical processing or analysis workflow for DTI that may result in much larger effects than shown in this work.

Eklund et al. (Eklund et al., 2016) used resting-state fMRI to obtain “null data”, i.e., truly negative data, to test the false-positive ratios for task fMRI. Unfortunately, for DTI, such an experimental testing setup to evaluate statistical inferences related to methodological factors is not trivial. However, without loss of generality, in this work, we performed a standard group study on gender as the framework to evaluate the effect of using different diffusion tensor estimation approaches. We used HCP data because of the excellent data quality and the large number of subjects with proper male-female balance, thereby eliminating issues related to small sample size and low power during statistical inference (Button et al., 2013).

In this work, we did not opt for analyzing the “statistical” significance (i.e., p-values) of our findings, but rather considered the difference in effect sizes that can be observed. In a similar context, shifting the focus from p-values to effect sizes was also recently presented by Ritchie et al. (Ritchie et al., 2018). They compared volumes and DTI based metrics of cortical, subcortical, and WM regions between females and males from the UK BioBank for more than 5000 participants. The comparison of the right CST revealed that males have larger FA values than females, with a p-value of 4×10^−65^ using Cohen’s d = 0.54. After adjusting for total brain volume, the values changed to 8×10^−12^ with Cohen’s d = 0.22. While these p-values are indeed *very* significant, they do not contain any useful information. On the other hand, the effect size measures provide more practical information. That is, adding another 5000 or more participants to the analysis will not result in any meaningful change in terms of the effect size, as this investigation is already statistically well-powered, while the p-value would decrease further. For the same reason, i.e., avoiding under-powered study design, we used HCP data for our group comparison, allowing us to focus on the performance of the DTI estimators.

Despite the efforts of optimizing the dMRI processing pipeline, it is often not clear what the benefits are of new developments for group-based studies. In this work, however, we showed that the application of IWLLS should be preferred over the OLLS for diffusion tensor estimation. The current framework can be easily extended to examine effects of modifying other processing elements, but also to investigate choices in algorithms and settings for specific analysis strategies, like tractography and connectomics, further improving the reliability and validity of future dMRI group studies.

## Supplementary Figures

**Suppl. Fig. 1.**
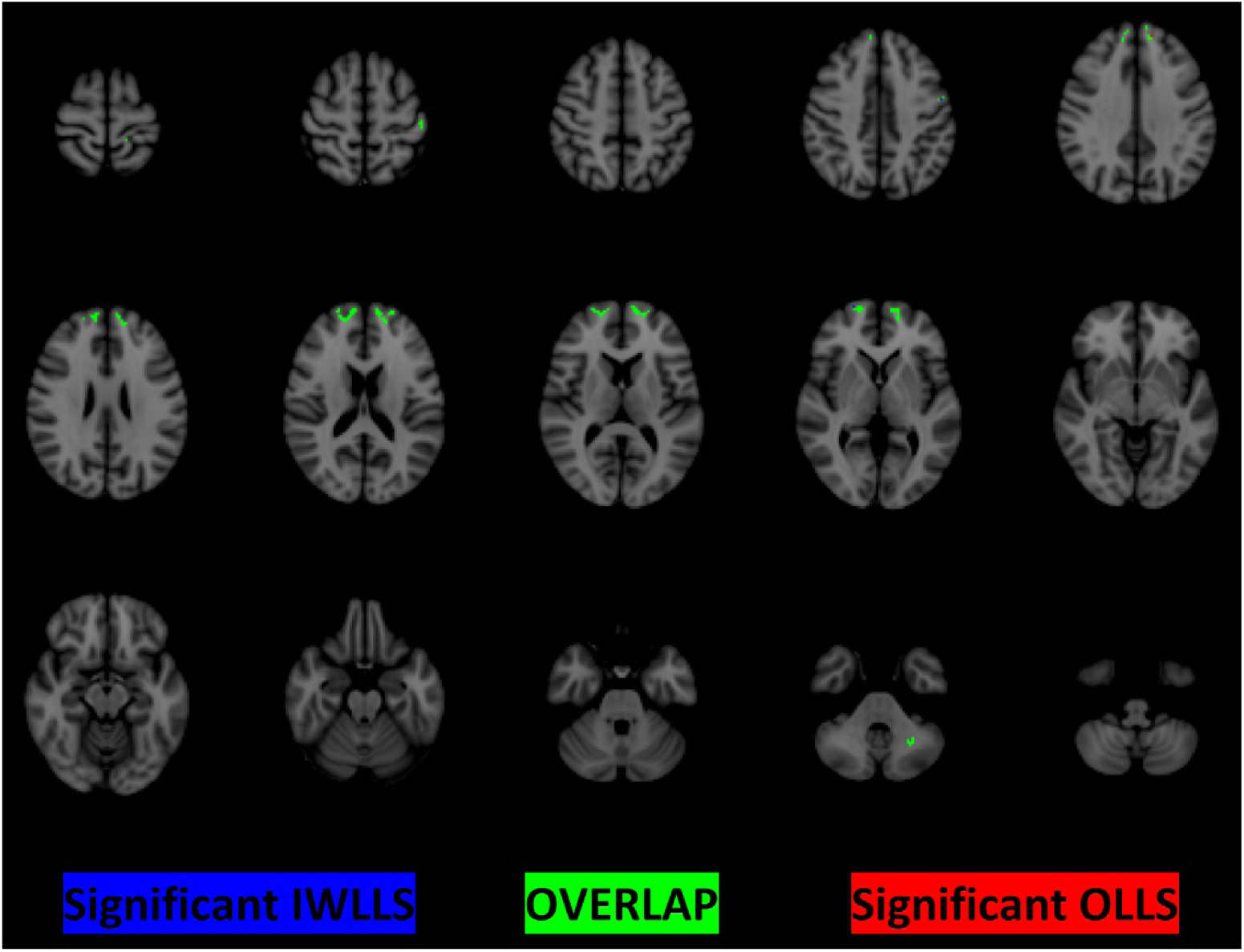
Results of the voxelwise analysis, indicating the regions where the FA is significantly higher for males than for females. Voxels colored in red and blue represent the regions where the FA estimates were obtained with the OLLS and IWLLS estimators, respectively (only visible in a few voxels). The green voxels show their overlap, i.e., the regions where both OLLS and IWLLS reflect significantly higher FA values for males compared to females.

## References

Andersson, J.L.R., 2008. Maximum a posteriori estimation of diffusion tensor parameters using a Rician noise model: Why, how and but. Neuroimage 42, 1340–1356. https://doi.org/10.1016/j.neuroimage.2008.05.053

Andersson, J.L.R., Graham, M.S., Drobnjak, I., Zhang, H., Campbell, J., 2018. Susceptibility-induced distortion that varies due to motion: Correction in diffusion MR without acquiring additional data. Neuroimage 171, 277–295. https://doi.org/10.1016/j.neuroimage.2017.12.040

Andersson, J.L.R., Graham, M.S., Zsoldos, E., Sotiropoulos, S.N., 2016. Incorporating outlier detection and replacement into a non-parametric framework for movement and distortion correction of diffusion MR images. Neuroimage 141, 556–572. https://doi.org/10.1016/j.neuroimage.2016.06.058

Andersson, J.L.R., Skare, S., Ashburner, J., 2003. How to correct susceptibility distortions in spin-echo echo-planar images: Application to diffusion tensor imaging. Neuroimage 20, 870–888. https://doi.org/10.1016/S1053-8119(03)00336-7

Andersson, J.L.R., Sotiropoulos, S.N., 2016. An integrated approach to correction for off-resonance effects and subject movement in diffusion MR imaging. Neuroimage 125, 1063–1078. https://doi.org/10.1016/j.neuroimage.2015.10.019

Andersson, J.L.R., Sotiropoulos, S.N., 2015. Non-parametric representation and prediction of single- and multi-shell diffusion-weighted MRI data using Gaussian processes. Neuroimage 122, 166–176. https://doi.org/10.1016/j.neuroimage.2015.07.067

Assaf, Y., Johansen-Berg, H., Thiebaut de Schotten, M., 2019. The role of diffusion MRI in neuroscience. NMR Biomed. 32, e3762. https://doi.org/10.1002/nbm.3762

Bach, M., Laun, F.B., Leemans, A., Tax, C.M.W., Biessels, G.J., Stieltjes, B., Maier-Hein, K.H., 2014. Methodological considerations on tract-based spatial statistics (TBSS). Neuroimage 100, 358–369. https://doi.org/10.1016/j.neuroimage.2014.06.021

Bammer, R., Markl, M., Barnett, A., Acar, B., Alley, M.T., Pelc, N.J., Glover, G.H., Moseley, M.E., 2003. Analysis and generalized correction of the effect of spatial gradient field distortions in diffusion-weighted imaging. Magn. Reson. Med. 50, 560–569. https://doi.org/10.1002/mrm.10545

Basser, P.J., Mattiello, J., LeBihan, D., 1994. MR diffusion tensor spectroscopy and imaging. Biophys. J. 66, 259–267. https://doi.org/10.1016/S0006-3495(94)80775-1

Bastiani, M., Cottaar, M., Fitzgibbon, S.P., Suri, S., Alfaro-Almagro, F., Sotiropoulos, S.N., Jbabdi, S., Andersson, J.L.R., 2019. Automated quality control for within and between studies diffusion MRI data using a non-parametric framework for movement and distortion correction. Neuroimage 184, 801–812. https://doi.org/10.1016/j.neuroimage.2018.09.073

Button, K.S., Ioannidis, J.P.A., Mokrysz, C., Nosek, B.A., Flint, J., Robinson, E.S.J., Munafò, M.R., 2013. Power failure: Why small sample size undermines the reliability of neuroscience. Nat. Rev. Neurosci. 14, 365–376. https://doi.org/10.1038/nrn3475

Caeyenberghs, K., Leemans, A., 2014. Hemispheric lateralization of topological organization in structural brain networks. Hum. Brain Mapp. 35, 4944–4957. https://doi.org/10.1002/hbm.22524

Catani, M., Thiebaut de Schotten, M., Slater, D., Dell’Acqua, F., 2013. Connectomic approaches before the connectome. Neuroimage 80, 2–13. https://doi.org/10.1016/j.neuroimage.2013.05.109

Cercignani, M., Gandini Wheeler-Kingshott, C., 2019. From micro-to macro-structures in multiple sclerosis: What is the added value of diffusion imaging. NMR Biomed. 32, 1–10. https://doi.org/10.1002/nbm.3888

Chang, L.C., Jones, D.K., Pierpaoli, C., 2005. RESTORE: Robust estimation of tensors by outlier rejection. Magn. Reson. Med. 53, 1088–1095. https://doi.org/10.1002/mrm.20426

Chang, L.C., Walker, L., Pierpaoli, C., 2012. Informed RESTORE: A method for robust estimation of diffusion tensor from low redundancy datasets in the presence of physiological noise artifacts. Magn. Reson. Med. 68, 1654–1663. https://doi.org/10.1002/mrm.24173

Collier, Q., Veraart, J., Jeurissen, B., Den Dekker, A.J., Sijbers, J., 2015. Iterative reweighted linear least squares for accurate, fast, and robust estimation of diffusion magnetic resonance parameters. Magn. Reson. Med. 73, 2174–2184. https://doi.org/10.1002/mrm.25351

Collier, Q., Veraart, J., Jeurissen, B., Vanhevel, F., Pullens, P., Parizel, P.M., den Dekker, A.J., Sijbers, J., 2018. Diffusion kurtosis imaging with free water elimination: A bayesian estimation approach. Magn. Reson. Med. 80, 802–813. https://doi.org/10.1002/mrm.27075

David, S., Heemskerk, A.M., Corrivetti, F., Thiebaut de Schotten, M., Sarubbo, S., Corsini, F., De Benedictis, A., Petit, L., Viergever, M.A., Jones, D.K., Mandonnet, E., Axer, H., Evans, J., Paus, T., Leemans, A., 2019. The Superoanterior Fasciculus (SAF): A Novel White Matter Pathway in the Human Brain? Front. Neuroanat. 13, 1–18. https://doi.org/10.3389/fnana.2019.00024

David, S., Tax, C.M.W., Viergever, M.A., Heemskerk, A.M., Leemans, A., 2015. Choices in processing steps for diffusion MRI analyses: Does it really matter?, in: Proceedings of the International Society for Magnetic Resonance in Medicine. p. 2981.

Eierud, C., Craddock, R.C., Fletcher, S., Aulakh, M., King-Casas, B., Kuehl, D., Laconte, S.M., 2014. Neuroimaging after mild traumatic brain injury: Review and meta-analysis. NeuroImage Clin. 4, 283–294. https://doi.org/10.1016/j.nicl.2013.12.009

Eklund, A., Nichols, T.E., Knutsson, H., 2016. Cluster failure: Why fMRI inferences for spatial extent have inflated false-positive rates. Proc. Natl. Acad. Sci. 113, 7900–7905. https://doi.org/10.1073/pnas.1602413113

ENIGMA DTI protocol, 2018. ENIGMA DTI protocol [WWW Document]. URL http://enigma.ini.usc.edu/protocols/dti-protocols/ (accessed 12.1.18).

Essen, D.C. Van, Ugurbil, K., Auerbach, E., Barch, D., Behrens, T.E.J.J., Bucholz, R., Chang, A., Chen, L., Corbetta, M., Curtiss, S.W., Penna, S. Della, Feinberg, D., Glasser, M.F., Harel, N., Heath, A.C., Larson-prior, L., Marcus, D., Michalareas, G., Moeller, S., Oostenveld, R., Petersen, S.E., Prior, F., Schlaggar, B.L., Smith, S.M., Snyder, A.Z., Xu, J., Yacoub, E., Consortium, W.H.C.P., Eeg, M.E.G., Van Essen, D.C., Ugurbil, K., Auerbach, E., Barch, D., Behrens, T.E.J.J., Bucholz, R., Chang, A., Chen, L., Corbetta, M., Curtiss, S.W., Della Penna, S., Feinberg, D., Glasser, M.F., Harel, N., Heath, A.C., Larson-prior, L., Marcus, D., Michalareas, G., Moeller, S., Oostenveld, R., Petersen, S.E., Prior, F., Schlaggar, B.L., Smith, S.M., Snyder, A.Z., Xu, J., Yacoub, E., 2012. The Human Connectome Project: A data acquisition perspective. Neuroimage 62, 2222–2231. https://doi.org/10.1016/j.neuroimage.2012.02.018

Filippini, N., Zsoldos, E., Haapakoski, R., Sexton, C.E., Mahmood, A., Allan, C.L., Topiwala, A., Valkanova, V., Brunner, E.J., Shipley, M.J., Auerbach, E., Moeller, S., Ugurbil, K., Xu, J., Yacoub, E., Andersson, J., Bijsterbosch, J., Clare, S., Griffanti, L., Hess, A.T., Jenkinson, M., Miller, K.L., Salimi-Khorshidi, G., Sotiropoulos, S.N., Voets, N.L., Smith, S.M., Geddes, J.R., Singh-Manoux, A., Mackay, C.E., Kivimäki, M., Ebmeier, K.P., 2014. Study protocol: The Whitehall II imaging sub-study. BMC Psychiatry 14, 159. https://doi.org/10.1186/1471-244X-14-159

Fonov, V., Evans, A.C., Botteron, K., Almli, C.R., McKinstry, R.C., Collins, D.L., 2011. Unbiased average age-appropriate atlases for pediatric studies. Neuroimage 54, 313–327. https://doi.org/10.1016/j.neuroimage.2010.07.033

Glasser, M.F., Sotiropoulos, S.N., Wilson, J.A., Coalson, T.S., Fischl, B., Andersson, J.L., Xu, J., Jbabdi, S., Webster, M., Polimeni, J.R., Van Essen, D.C., Jenkinson, M., 2013. The minimal preprocessing pipelines for the Human Connectome Project. Neuroimage 80, 105–124. https://doi.org/10.1016/j.neuroimage.2013.04.127

Graham, M.S., Drobnjak, I., Jenkinson, M., Zhang, H., 2017. Quantitative assessment of the susceptibility artefact and its interaction with motion in diffusion MRI. PLoS One 12, 1–25. https://doi.org/10.1371/journal.pone.0185647

HCP, 2017. HCP WIKI [WWW Document]. URL https://wiki.humanconnectome.org/display/PublicData/HCP+Data+Release+Updates%3A+Known+Issues+and+Planned+fixes (accessed 5.10.18).

Herting, M.M., Maxwell, E.C., Irvine, C., Nagel, B.J., 2012. The impact of sex, puberty, and hormones on white matter microstructure in adolescents. Cereb. Cortex 22, 1979–1992. https://doi.org/10.1093/cercor/bhr246

Holmes, A.P., Blair, R.C., Watson, &NA; G., Ford, I., Watson, J.D.G.G., Ford, I., Watson, H.J.D.G., Ford, I., 1996. Nonparametric Analysis of Statistic Images from Functional Mapping Experiments. J. Cereb. Blood Flow Metab. 16, 7–22. https://doi.org/10.1097/00004647-199601000-00002

Hsu, J.-L.L., Leemans, A., Bai, C.-H.H., Lee, C.-H.H., Tsai, Y.-F.F., Chiu, H.-C.C., Chen, W.-H.H., 2008. Gender differences and age-related white matter changes of the human brain: A diffusion tensor imaging study. Neuroimage 39, 566–577. https://doi.org/10.1016/j.neuroimage.2007.09.017

Ingalhalikar, M., Smith, A., Parker, D., Satterthwaite, T.D., Elliott, M.A., Ruparel, K., Hakonarson, H., Gur, R.E., Gur, R.C., Verma, R., 2014. Sex differences in the structural connectome of the human brain. Proc. Natl. Acad. Sci. 111, 823–828. https://doi.org/10.1073/pnas.1316909110

Jenkinson, M., Beckmann, C.F., Behrens, T.E.J., Woolrich, M.W., Smith, S.M., 2012. Fsl. Neuroimage 62, 782–790. https://doi.org/10.1016/j.neuroimage.2011.09.015

Jones, D.K., Basser, P.J., 2004. “Squashing peanuts and smashing pumpkins”: How noise distorts diffusion-weighted MR data. Magn. Reson. Med. 52, 979–993. https://doi.org/10.1002/mrm.20283

Kanaan, R.A., Allin, M., Picchioni, M., Barker, G.J., Daly, E., Shergill, S.S., Woolley, J., McGuire, P.K., 2012. Gender differences in white matter microstructure. PLoS One 7. https://doi.org/10.1371/journal.pone.0038272

Kellner, E., Dhital, B., Kiselev, V.G., Reisert, M., 2016. Gibbs-ringing artifact removal based on local subvoxel-shifts. Magn. Reson. Med. 76, 1574–1581. https://doi.org/10.1002/mrm.26054

Koay, C.G., Özarslan, E., Basser, P.J., 2009. A signal transformational framework for breaking the noise floor and its applications in MRI. J. Magn. Reson. 197, 108–119. https://doi.org/10.1016/j.jmr.2008.11.015

Kozák, L.R., David, S., Rudas, G., Vidnyánszky, Z., Leemans, A., Nagy, Z., 2013. Investigating the need of triggering the acquisition for infant diffusion MRI: A quantitative study including bootstrap statistics. Neuroimage 69, 198–205. https://doi.org/10.1016/j.neuroimage.2012.11.063

Kristoffersen, A., 2012. Estimating non-Gaussian diffusion model parameters in the presence of physiological noise and Rician signal bias. J. Magn. Reson. Imaging 35, 181–189. https://doi.org/10.1002/jmri.22826

Kristoffersen, A., 2007. Optimal estimation of the diffusion coefficient from non-averaged and averaged noisy magnitude data. J. Magn. Reson. 187, 293–305. https://doi.org/10.1016/j.jmr.2007.05.004

Leemans, A., Jeurissen, B., Sijbers, J., Jones, D.K., Jeruissen, B., Sijbers, J., Jones, D.K., 2009. ExploreDTI: a graphical toolbox for processing, analyzing, and visualizing diffusion MR data. Proc. Int. Soc. Magn. Reson. Med. 17, 3537. https://doi.org/10.1093/occmed/kqr069

Leemans, A., Jones, D.K., 2009. The B-matrix must be rotated when correcting for subject motion in DTI data. Magn. Reson. Med. 61, 1336–1349. https://doi.org/10.1002/mrm.21890

Lunven, M., De Schotten, M.T., Bourlon, C., Duret, C., Migliaccio, R., Rode, G., Bartolomeo, P., 2015. White matter lesional predictors of chronic visual neglect: A longitudinal study. Brain 138, 746–760. https://doi.org/10.1093/brain/awu389

Lustig, M., Donoho, D., Pauly, J.M., 2007. Sparse MRI: The application of compressed sensing for rapid MR imaging. Magn. Reson. Med. 58, 1182–1195. https://doi.org/10.1002/mrm.21391

McNab, J.A., Edlow, B.L., Witzel, T., Huang, S.Y., Bhat, H., Heberlein, K., Feiweier, T., Liu, K., Keil, B., Cohen-Adad, J., Tisdall, M.D., Folkerth, R.D., Kinney, H.C., Wald, L.L., 2013. The Human Connectome Project and beyond: Initial applications of 300mT/m gradients. Neuroimage 80, 234–245. https://doi.org/10.1016/j.neuroimage.2013.05.074

Menzler, K., Belke, M., Wehrmann, E., Krakow, K., Lengler, U., Jansen, A., Hamer, H.M., Oertel, W.H., Rosenow, F., Knake, S., 2011. Men and women are different: Diffusion tensor imaging reveals sexual dimorphism in the microstructure of the thalamus, corpus callosum and cingulum. Neuroimage 54, 2557–2562. https://doi.org/10.1016/j.neuroimage.2010.11.029

Mesri, H.Y., David, S., Viergever, M.A., Leemans, A., 2019. The adverse effect of gradient nonlinearities on diffusion MRI: From voxels to group studies. Neuroimage 116127. https://doi.org/10.1016/J.NEUROIMAGE.2019.116127

Moser, E., Laistler, E., Schmitt, F., Kontaxis, G., 2017. High Field NMR and MRI–The Role of Magnet Technology to Increase Sensitivity and Specificity. Front. Phys. 5, 33. https://doi.org/10.3389/fphy.2017.00041

Mueller, S.G., Weiner, M.W., Thal, L.J., Petersen, R.C., Jack, C., Jagust, W., Trojanowski, J.Q., Toga, A.W., Beckett, L., 2005. The Alzheimer’s disease neuroimaging initiative. Neuroimaging Clin. N. Am. 15, 869–877. https://doi.org/10.1016/j.nic.2005.09.008

Nichols, T., Holmes, A., 2003. Nonparametric Permutation Tests for Functional Neuroimaging. Hum. Brain Funct. Second Ed. 25, 887–910. https://doi.org/10.1016/B978-012264841-0/50048-2

Nir, T.M., Jahanshad, N., Villalon-Reina, J.E., Toga, A.W., Jack, C.R., Weiner, M.W., Thompson, P.M., 2013. Effectiveness of regional DTI measures in distinguishing Alzheimer’s disease, MCI, and normal aging. NeuroImage Clin. 3, 180–195. https://doi.org/10.1016/j.nicl.2013.07.006

Nosek, B.A., Lakens, D., 2014. Registered reports: A method to increase the credibility of published results. Soc. Psychol. (Gott). 45, 137–141. https://doi.org/10.1027/1864-9335/a000192

Novikov, D.S., Fieremans, E., Jespersen, S.N., Kiselev, V.G., 2019. Quantifying brain microstructure with diffusion MRI: Theory and parameter estimation. NMR Biomed. 32, e3998. https://doi.org/10.1002/nbm.3998

Núžez, C., Theofanopoulou, C., Senior, C., Cambra, M.R., Usall, J., Stephan-otto, C., Brébion, G., Nu, C., Cambra, M.R., Usall, J., Stephan-otto, C., Bre, G., 2017. A large-scale study on the effects of sex on gray matter asymmetry. Brain Struct. Funct. 1–11. https://doi.org/10.1007/s00429-017-1481-4

Owen, J.P., Marco, E.J., Desai, S., Fourie, E., Harris, J., Hill, S.S., Arnett, A.B., Mukherjee, P., 2013. Abnormal white matter microstructure in children with sensory processing disorders. NeuroImage Clin. 2, 844–853. https://doi.org/10.1016/j.nicl.2013.06.009

Pannek, K., Raffelt, D., Bell, C., Mathias, J.L., Rose, S.E., 2012. HOMOR: Higher Order Model Outlier Rejection for high b-value MR diffusion data. Neuroimage 63, 835–842. https://doi.org/10.1016/j.neuroimage.2012.07.022

Penny, W., Friston, K., Ashburner, J., Kiebel, S., Nichols, T., 2007. Statistical Parametric Mapping: The Analysis of Functional Brain Images. Elsevier. https://doi.org/10.1016/B978-0-12-372560-8.X5000-1

Perrone, D., Aelterman, J., Pižurica, A., Jeurissen, B., Philips, W., Leemans, A., 2015. The effect of Gibbs ringing artifacts on measures derived from diffusion MRI. Neuroimage 120, 441–455. https://doi.org/10.1016/j.neuroimage.2015.06.068

Phillips, O.R., Joshi, S.H., Piras, F., Orfei, M.D., Iorio, M., Narr, K.L., Shattuck, D.W., Caltagirone, C., Spalletta, G., Di Paola, M., 2016. The superficial white matter in Alzheimer’s disease. Hum. Brain Mapp. 37, 1321–1334. https://doi.org/10.1002/hbm.23105

Poldrack, R.A., 2012. The future of fMRI in cognitive neuroscience. Neuroimage 62, 1216–1220. https://doi.org/10.1016/j.neuroimage.2011.08.007

Poldrack, R.A., Baker, C.I., Durnez, J., Gorgolewski, K.J., Matthews, P.M., Munafò, M.R., Nichols, T.E., Poline, J.B., Vul, E., Yarkoni, T., 2017. Scanning the horizon: Towards transparent and reproducible neuroimaging research. Nat. Rev. Neurosci. 18, 115–126. https://doi.org/10.1038/nrn.2016.167

Ritchie, S.J., Cox, S.R., Shen, X., Lombardo, M. V, Reus, L.M., Alloza, C., Harris, M.A., Alderson, H.L., Hunter, S., Neilson, E., Liewald, D.C.M., Auyeung, B., Whalley, H.C., Lawrie, S.M., Gale, C.R., Bastin, M.E., McIntosh, A.M., Deary, I.J., 2018. Sex Differences in the Adult Human Brain: Evidence from 5216 UK Biobank Participants. Cereb. Cortex 28, 2959–2975. https://doi.org/10.1093/cercor/bhy109

Rothstein, H.R., Sutton, A.J., Borenstein, M., 2006. Publication Bias in Meta-Analysis: Prevention, Assessment and Adjustments, Publication Bias in Meta-Analysis: Prevention, Assessment and Adjustments. Wiley. https://doi.org/10.1002/0470870168

Rousselet, G.A., Pernet, C.R., Wilcox, R.R., 2017. Beyond differences in means: robust graphical methods to compare two groups in neuroscience. Eur. J. Neurosci. 46, 1738–1748. https://doi.org/10.1111/ejn.13610

Rudie, J.D., Brown, J.A., Beck-Pancer, D., Hernandez, L.M., Dennis, E.L., Thompson, P.M., Bookheimer, S.Y., Dapretto, M., 2013. Altered functional and structural brain network organization in autism. NeuroImage Clin. 2, 79–94. https://doi.org/10.1016/j.nicl.2012.11.006

Sabia, S., Dugravot, A., Dartigues, J.F., Abell, J., Elbaz, A., Kivimäki, M., Singh-Manoux, A., 2017. Physical activity, cognitive decline, and risk of dementia: 28 year follow-up of Whitehall II cohort study. BMJ 357, j2709. https://doi.org/10.1136/bmj.j2709

Salvador, R., Peža, A., Menon, D.K., Carpenter, T.A., Pickard, J.D., Bullmore, E.T., 2005. Formal characterization and extension of the linearized diffusion tensor model. Hum. Brain Mapp. 24, 144–155. https://doi.org/10.1002/hbm.20076

Schwarz, S.T., Abaei, M., Gontu, V., Morgan, P.S., Bajaj, N., Auer, D.P., 2013. Diffusion tensor imaging of nigral degeneration in Parkinson’s disease: A region-of-interest and voxel-based study at 3 T and systematic review with meta-analysis. NeuroImage Clin. 3, 481–488. https://doi.org/10.1016/j.nicl.2013.10.006

Setsompop, K., Kimmlingen, R., Eberlein, E., Witzel, T., Cohen-Adad, J., McNab, J.A., Keil, B., Tisdall, M.D., Hoecht, P., Dietz, P., Cauley, S.F., Tountcheva, V., Matschl, V., Lenz, V.H., Heberlein, K., Potthast, A., Thein, H., Van Horn, J., Toga, A., Schmitt, F., Lehne, D., Rosen, B.R., Wedeen, V., Wald, L.L., 2013. Pushing the limits of in vivo diffusion MRI for the Human Connectome Project. Neuroimage 80, 220–233. https://doi.org/10.1016/j.neuroimage.2013.05.078

Smaldino, P.E., McElreath, R., 2016. the Natural Selection of Bad Science. R. Soc. Open Sci. 3, 1–20. https://doi.org/10.1098/rsos.160384

Smith, S.M., Jenkinson, M., Johansen-Berg, H., Rueckert, D., Nichols, T.E., Mackay, C.E., Watkins, K.E., Ciccarelli, O., Cader, M.Z., Matthews, P.M., Behrens, T.E.J., 2006. Tract-based spatial statistics: Voxelwise analysis of multi-subject diffusion data. Neuroimage 31, 1487–1505. https://doi.org/10.1016/j.neuroimage.2006.02.024

Smith, S.M., Nichols, T.E., 2018. Statistical Challenges in “Big Data” Human Neuroimaging. Neuron 97, 263–268. https://doi.org/10.1016/j.neuron.2017.12.018

Smith, S.M., Nichols, T.E., 2009. Threshold-free cluster enhancement: Addressing problems of smoothing, threshold dependence and localisation in cluster inference. Neuroimage 44, 83–98. https://doi.org/10.1016/j.neuroimage.2008.03.061

Sotiropoulos, S.N., Jbabdi, S., Xu, J., Andersson, J.L., Moeller, S., Auerbach, E.J., Glasser, M.F., Hernandez, M., Sapiro, G., Jenkinson, M., Feinberg, D.A., Yacoub, E., Lenglet, C., Van Essen, D.C., Ugurbil, K., Behrens, T.E.J., 2013. Advances in diffusion MRI acquisition and processing in the Human Connectome Project. Neuroimage 80, 125–143. https://doi.org/10.1016/j.neuroimage.2013.05.057

St-Jean, S., Coupé, P., Descoteaux, M., 2016. Non Local Spatial and Angular Matching: Enabling higher spatial resolution diffusion MRI datasets through adaptive denoising. Med. Image Anal. 32, 115–130. https://doi.org/10.1016/j.media.2016.02.010

Sudlow, C., Gallacher, J., Allen, N., Beral, V., Burton, P., Danesh, J., Downey, P., Elliott, P., Green, J., Landray, M., Liu, B., Matthews, P., Ong, G., Pell, J., Silman, A., Young, A., Sprosen, T., Peakman, T., Collins, R., 2015. UK Biobank: An Open Access Resource for Identifying the Causes of a Wide Range of Complex Diseases of Middle and Old Age. PLoS Med. 12, 1001779. https://doi.org/10.1371/journal.pmed.1001779

Tax, C.M.W., Otte, W.M., Viergever, M.A., Dijkhuizen, R.M., Leemans, A., 2015. REKINDLE: Robust Extraction of Kurtosis INDices with Linear Estimation. Magn. Reson. Med. 73, 794–808. https://doi.org/10.1002/mrm.25165

Thiebaut de Schotten, M., Dell’Acqua, F., Valabregue, R., Catani, M., 2012. Monkey to human comparative anatomy of the frontal lobe association tracts. Cortex 48, 82–96. https://doi.org/10.1016/j.cortex.2011.10.001

Thompson, P.M., Hibar, D.P., Stein, J.L., Prasad, G., Jahanshad, N., 2016. Genetics of the connectome and the ENIGMA project, in: Research and Perspectives in Neurosciences. Springer, Cham, pp. 147–164. https://doi.org/10.1007/978-3-319-27777-6_10

Thompson, P.M., Stein, J.L., Medland, S.E., Hibar, D.P., Vasquez, A.A., Renteria, M.E., Toro, R., Jahanshad, N., Schumann, G., Franke, B., Wright, M.J., Martin, N.G., Agartz, I., Alda, M., Alhusaini, S., Almasy, L., Almeida, J., Alpert, K., Andreasen, N.C., Andreassen, O.A., Apostolova, L.G., Appel, K., Armstrong, N.J., Aribisala, B., Bastin, M.E., Bauer, M., Bearden, C.E., Bergmann, Ø., Binder, E.B., Blangero, J., Bockholt, H.J., Bøen, E., Bois, C., Boomsma, D.I., Booth, T., Bowman, I.J., Bralten, J., Brouwer, R.M., Brunner, H.G., Brohawn, D.G., Buckner, R.L., Buitelaar, J., Bulayeva, K., Bustillo, J.R., Calhoun, V.D., Cannon, D.M., Cantor, R.M., Carless, M.A., Caseras, X., Cavalleri, G.L., Chakravarty, M.M., Chang, K.D., Ching, C.R.K., Christoforou, A., Cichon, S., Clark, V.P., Conrod, P., Coppola, G., Crespo-Facorro, B., Curran, J.E., Czisch, M., Deary, I.J., de Geus, E.J.C., den Braber, A., Delvecchio, G., Depondt, C., de Haan, L., de Zubicaray, G.I., Dima, D., Dimitrova, R., Djurovic, S., Dong, H., Donohoe, G., Duggirala, R., Dyer, T.D., Ehrlich, S., Ekman, C.J., Elvsåshagen, T., Emsell, L., Erk, S., Espeseth, T., Fagerness, J., Fears, S., Fedko, I., Fernández, G., Fisher, S.E., Foroud, T., Fox, P.T., Francks, C., Frangou, S., Frey, E.M., Frodl, T., Frouin, V., Garavan, H., Giddaluru, S., Glahn, D.C., Godlewska, B., Goldstein, R.Z., Gollub, R.L., Grabe, H.J., Grimm, O., Gruber, O., Guadalupe, T., Gur, R.E., Gur, R.C., Göring, H.H.H., Hagenaars, S., Hajek, T., Hall, G.B., Hall, J., Hardy, J., Hartman, C.A., Hass, J., Hatton, S.N., Haukvik, U.K., Hegenscheid, K., Heinz, A., Hickie, I.B., Ho, B.C., Hoehn, D., Hoekstra, P.J., Hollinshead, M., Holmes, A.J., Homuth, G., Hoogman, M., Hong, L.E., Hosten, N., Hottenga, J.J., Hulshoff Pol, H.E., Hwang, K.S., Jack, C.R., Jenkinson, M., Johnston, C., Jönsson, E.G., Kahn, R.S., Kasperaviciute, D., Kelly, S., Kim, S., Kochunov, P., Koenders, L., Krämer, B., Kwok, J.B.J., Lagopoulos, J., Laje, G., Landen, M., Landman, B.A., Lauriello, J., Lawrie, S.M., Lee, P.H., Le Hellard, S., Lemaître, H., Leonardo, C.D., Li, C. shan, Liberg, B., Liewald, D.C., Liu, X., Lopez, L.M., Loth, E., Lourdusamy, A., Luciano, M., Macciardi, F., Machielsen, M.W.J., MacQueen, G.M., Malt, U.F., Mandl, R., Manoach, D.S., Martinot, J.L., Matarin, M., Mather, K.A., Mattheisen, M., Mattingsdal, M., Meyer-Lindenberg, A., McDonald, C., McIntosh, A.M., McMahon, F.J., McMahon, K.L., Meisenzahl, E., Melle, I., Milaneschi, Y., Mohnke, S., Montgomery, G.W., Morris, D.W., Moses, E.K., Mueller, B.A., Mužoz Maniega, S., Mühleisen, T.W., Müller-Myhsok, B., Mwangi, B., Nauck, M., Nho, K., Nichols, T.E., Nilsson, L.G., Nugent, A.C., Nyberg, L., Olvera, R.L., Oosterlaan, J., Ophoff, R.A., Pandolfo, M., Papalampropoulou-Tsiridou, M., Papmeyer, M., Paus, T., Pausova, Z., Pearlson, G.D., Penninx, B.W., Peterson, C.P., Pfennig, A., Phillips, M., Pike, G.B., Poline, J.B., Potkin, S.G., Pütz, B., Ramasamy, A., Rasmussen, J., Rietschel, M., Rijpkema, M., Risacher, S.L., Roffman, J.L., Roiz-Santiažez, R., Romanczuk-Seiferth, N., Rose, E.J., Royle, N.A., Rujescu, D., Ryten, M., Sachdev, P.S., Salami, A., Satterthwaite, T.D., Savitz, J., Saykin, A.J., Scanlon, C., Schmaal, L., Schnack, H.G., Schork, A.J., Schulz, S.C., Schür, R., Seidman, L., Shen, L., Shoemaker, J.M., Simmons, A., Sisodiya, S.M., Smith, C., Smoller, J.W., Soares, J.C., Sponheim, S.R., Sprooten, E., Starr, J.M., Steen, V.M., Strakowski, S., Strike, L., Sussmann, J., Sämann, P.G., Teumer, A., Toga, A.W., Tordesillas-Gutierrez, D., Trabzuni, D., Trost, S., Turner, J., Van den Heuvel, M., van der Wee, N.J., van Eijk, K., van Erp, T.G.M., van Haren, N.E.M., van’t Ent, D., van Tol, M.J., Valdés Hernández, M.C., Veltman, D.J., Versace, A., Völzke, H., Walker, R., Walter, H., Wang, L., Wardlaw, J.M., Weale, M.E., Weiner, M.W., Wen, W., Westlye, L.T., Whalley, H.C., Whelan, C.D., White, T., Winkler, A.M., Wittfeld, K., Woldehawariat, G., Wolf, C., Zilles, D., Zwiers, M.P., Thalamuthu, A., Schofield, P.R., Freimer, N.B., Lawrence, N.S., Drevets, W., 2014. The ENIGMA Consortium: Large-scale collaborative analyses of neuroimaging and genetic data. Brain Imaging Behav. 8, 153–182. https://doi.org/10.1007/s11682-013-9269-5

Tyan, Y.S., Liao, J.R., Shen, C.Y., Lin, Y.C., Weng, J.C., 2017. Gender differences in the structural connectome of the teenage brain revealed by generalized q-sampling MRI. NeuroImage Clin. 15, 376–382. https://doi.org/10.1016/j.nicl.2017.05.014

Veraart, J., Fieremans, E., Jelescu, I.O., Knoll, F., Novikov, D.S., 2016a. Gibbs ringing in diffusion MRI. Magn. Reson. Med. 76, 301–314. https://doi.org/10.1002/mrm.25866

Veraart, J., Hecke, W. Van, Sijbers, J., Van Hecke, W., Sijbers, J., 2011. Constrained maximum likelihood estimation of the diffusion kurtosis tensor using a Rician noise model. Magn. Reson. Med. 66, 678–686. https://doi.org/10.1002/mrm.22835

Veraart, J., Novikov, D.S., Christiaens, D., Ades-aron, B., Sijbers, J., Fieremans, E., 2016b. Denoising of diffusion MRI using random matrix theory. Neuroimage 142, 394–406. https://doi.org/10.1016/j.neuroimage.2016.08.016

Veraart, J., Rajan, J., Peeters, R.R., Leemans, A., Sunaert, S., Sijbers, J., 2013a. Comprehensive framework for accurate diffusion MRI parameter estimation. Magn. Reson. Med. 70, 972–984. https://doi.org/10.1002/mrm.24529

Veraart, J., Sijbers, J., Sunaert, S., Leemans, A., Jeurissen, B., 2013b. Weighted linear least squares estimation of diffusion MRI parameters: Strengths, limitations, and pitfalls. Neuroimage 81, 335–346. https://doi.org/10.1016/j.neuroimage.2013.05.028

Vos, S.B., Tax, C.M.W., Luijten, P.R., Ourselin, S., Leemans, A., Froeling, M., 2017. The importance of correcting for signal drift in diffusion MRI. Magn. Reson. Med. 77, 285–299. https://doi.org/10.1002/mrm.26124

Wasserstein, R.L., Lazar, N.A., 2016. The ASA’s Statement on *p* -Values: Context, Process, and Purpose. Am. Stat. 70, 129–133. https://doi.org/10.1080/00031305.2016.1154108

Westerhausen, R., Walter, C., Kreuder, F., Wittling, R.A., Schweiger, E., Wittling, W., 2003. The influence of handedness and gender on the microstructure of the human corpus callosum: A diffusion-tensor magnetic resonance imaging study. Neurosci. Lett. 351, 99–102. https://doi.org/10.1016/j.neulet.2003.07.011

Wierenga, L.M., Sexton, J.A., Laake, P., Giedd, J.N., Tamnes, C.K., 2017. A Key Characteristic of Sex Differences in the Developing Brain: Greater Variability in Brain Structure of Boys than Girls. Cereb. Cortex 1–11. https://doi.org/10.1093/cercor/bhx154

Wilcox, R.R., 2012. Introduction to robust estimation and hypothesis testing. Academic Press.

Winkler, A.M., Ridgway, G.R., Douaud, G., Nichols, T.E., Smith, S.M., 2016. Faster permutation inference in brain imaging. Neuroimage 141, 502–516. https://doi.org/10.1016/j.neuroimage.2016.05.068

Winkler, A.M., Ridgway, G.R., Webster, M.A., Smith, S.M., Nichols, T.E., 2014. Permutation inference for the general linear model. Neuroimage 92, 381–397. https://doi.org/10.1016/j.neuroimage.2014.01.060

